# Firing Rate-dependent Phase Responses of Purkinje Cells Support Transient Oscillations

**DOI:** 10.1101/822478

**Authors:** Yunliang Zang, Sungho Hong, Erik De Schutter

## Abstract

Both spike rate and timing can transmit information in the brain. Phase response curves (PRCs) quantify how a neuron transforms input to output by spike timing. They exhibit strong firing-rate adaptation, but its mechanism and relevance for network output are poorly understood. Using our Purkinje cell (PC) model we demonstrate that the rate adaptation is caused by rate-dependent subthreshold membrane potentials efficiently regulating the activation of Na^+^ channels. Then we use a realistic PC network model to examine how rate-dependent responses synchronize spikes in the scenario of reciprocal inhibition-caused high-frequency oscillations. The changes in PRC cause oscillations and spike correlations only at high firing rates. The causal role of the PRC is confirmed using a simpler coupled oscillator network model. This mechanism enables transient oscillations between fast-spiking neurons that thereby form PC assemblies. Our work demonstrates that rate adaptation of PRCs can spatio-temporally organize the PC input to cerebellar nuclei.

## Introduction

The propensity of neurons to fire synchronously depends on the interaction between cellular and network properties (Ermentrout et al., 2001). The contribution of cellular properties can be measured with a phase response curve (PRC). The PRC quantifies how a weak stimulus exerted at different phases during the interspike interval (ISI) can shift subsequent spike timing in repetitively firing neurons (Ermentrout et al., 2001; Gutkin et al., 2005) and thereby predicts how well-timed synaptic input can modify spike timing. Consequently, the PRC determines the potential of network synchronization (Ermentrout et al., 2001; Ermentrout et al., 2008; Gutkin et al., 2005; Smeal et al., 2010). However, it is not static and shows significant adaptation to firing rates. In cerebellar Purkinje Cells (PCs), their phase responses to weak stimuli at low firing rates are small and surprisingly flat. With increased rates, responses in later phases become phase-dependent, with earlier onset-phases and gradually increasing peak amplitudes. This PRC property has never been theoretically replicated or explained (Couto et al., 2015; Phoka et al., 2010), nor has its effect on synchronizing spike outputs been explored.

On the circuit level, high frequency oscillations caused by reciprocal inhibition have been observed in many regions of the brain, including cortex, cerebellum and hippocampus (Bartos et al., 2002; Buzsaki and Draguhn, 2004; Cheron et al., 2004; de Solages et al., 2008). The functional importance of oscillations in information transmission is largely determined by their spatio-temporal scale, which for hard-wired inhibitory connections, is generally assumed to be driven by external input. It is interesting to explore whether firing rate-dependent PRCs can contribute to dynamic control of the spatial range of oscillations based on firing rate changes, because this would have significant downstream effects (Person and Raman, 2012).

To examine the mechanism of rate-dependent PRCs, we use our physiologically detailed PC model (Zang et al., 2018) and a simple pyramidal neuron model to explore the rate adaptation of PRCs. By analyzing simulation data and *in vitro* experimental data (Rancz and Hausser, 2010), we show that rate-dependent subthreshold membrane potentials can modulate the activation of Na^+^ channels to shape neuronal PRC profiles. We also build a PC network model connected by inhibitory axon collaterals to simulate high-frequency oscillations (de Solages et al., 2008; Witter et al., 2016). Rate adaptation of PRCs increases the power of oscillations at higher firing rates, firing irregularity and network connectivity also regulate the oscillation level. The causal role of the PRC is confirmed using a simpler coupled oscillator network model. The combination of these factors enables PC spikes uncorrelated at low basal rates to become transiently correlated in transient assemblies of PCs at high firing rates.

## Results

### PRC Exhibits Rate Adaptation in PCs

PRCs were obtained by repeatedly exerting a weak stimulus at different phases of the ISI. The resulting change in ISI relative to original ISI corresponds to the PRC value at that phase (Fig. 1A). All previous abstract and detailed PC models failed to replicate the experimentally observed rate adaptation of PRCs (Akemann and Knopfel, 2006; Couto et al., 2015; De Schutter and Bower, 1994; Khaliq et al., 2003; Phoka et al., 2010). Our recent PC model was well constrained against a wide range of experimental data (Zang et al., 2018). Here, we explored whether this model can capture the rate adaptation of PRCs under similar conditions. When the PC model fires at 12 Hz, responses (phase advances) to weak stimuli are small and nearly flat for the whole ISI (Fig. 1B, D). Only at a very narrow late phase do the responses become phase-dependent and slightly increased. With increased rates, the responses remain small and flat during early phases. However, later phase-dependent peaks gradually become larger (Fig. 1C), with onset shifted to earlier phases (Fig. 1D). It should be noted that the increased late-peak amplitude may be affected by how the PRC is computed (Eq. 1): it is normalized by the ISI, causing the peak amplitude to increase for higher firing rates (smaller ISIs).

**Figure 1.**
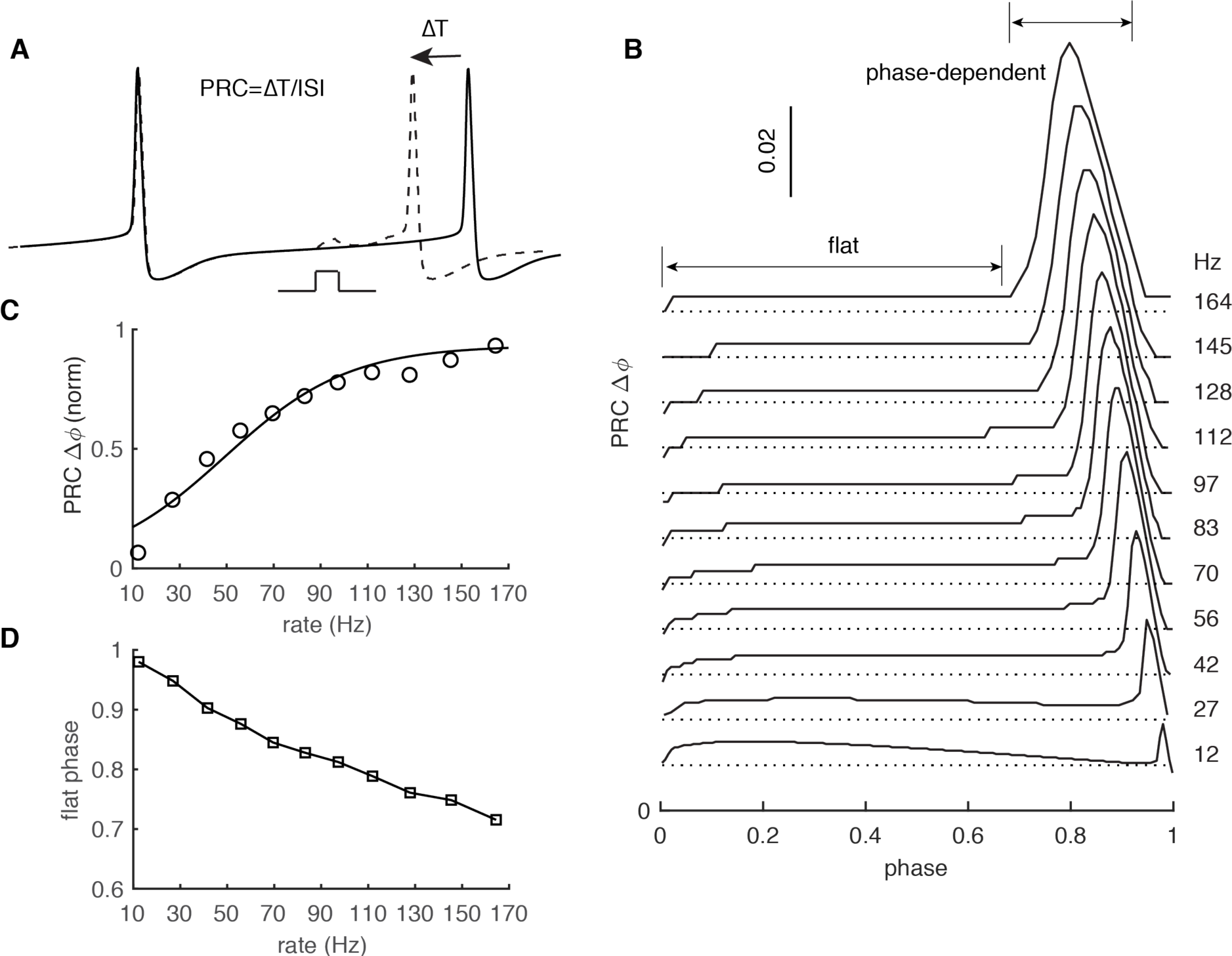
PRC Exhibits Strong Rate Adaptation in PC model. (**A**) Schematic representation of the definition and computation of PRCs. The current pulse has a duration of 0.5 ms and an amplitude of 50 pA. Different spike rates were achieved by somatic current injection (Couto et al., 2015; Phoka et al., 2010). (**B)** The rate adaptation of the flat part and the phase-dependent PRC peak. (**C)** PRC peak amplitudes at different firing rates fitted by the Boltzmann function. (**D**) Duration of the flat phase at different firing rates.

In agreement with experiments under the same stimulus conditions (Phoka et al., 2010), the peak of PRCs finally became saturated at ∼ 0.06 at high rates. The relationship between normalized PRC peaks and rates can be fitted by the Boltzman function and matches experimental data (Fig. 1C, fitted with 1/(1 + e^−(rate−a)/b^), a = 49.1, b = 26.4 in the model versus a = 44.1 and b = 20.5 in experiments (Couto et al., 2015). PRCs in our model show similar rate adaption with inhibitory stimuli (phase delay, Fig. S1A). This form of rate adaptive PRCs requires the presence of a dendrite in the PC model (Fig. S2), but the dendrite can be passive (Fig. S1B). We also tested the effect of increasing stimulus amplitude on PRC adaptation. Increasing stimulus amplitude consistently shifts onset-phases of phase-dependent peaks to the left and increases their amplitudes (Fig. S1C).

To unveil the biophysical principles governing rate adaptive PRC profiles, we need to answer two questions: why are responses flat in early phases and why do responses become phase-dependent during later phases?

### The Biophysical Mechanism of Rate Adaptation of PRCs in PCs

We examined how spike properties vary with firing rates and find that the facilitation of Na^+^ currents relative to K^+^ currents, due to elevated subthreshold membrane potentials at high rates, underlies the rate adaptation of PRCs. After each spike, there is a pronounced after-hyperpolarization (AHP) caused by the large conductance Ca^2+^-activated K^+^ current, and subsequently the membrane potential gradually depolarizes due to intrinsic Na^+^ currents and dendritic axial current (Zang et al., 2018). As confirmed by re-analyzing *in vitro* somatic membrane potential recordings (shared by Ede Rancz and Michael Häusser (Rancz and Hausser, 2010)), subthreshold membrane potential levels are significantly elevated at high firing rates, but spike thresholds rise only slightly with rates (Fig. 2A). This means that the ISI phase where Na^+^ activation threshold (∼ -55 mV for 0.5% activation in PCs (Khaliq et al., 2003; Zang et al., 2018)) is crossed shifts to earlier phases with increasing rates. Consequently, larger phase ranges of membrane potentials are above the threshold at high rates (Fig. 2B).

**Figure 2.**
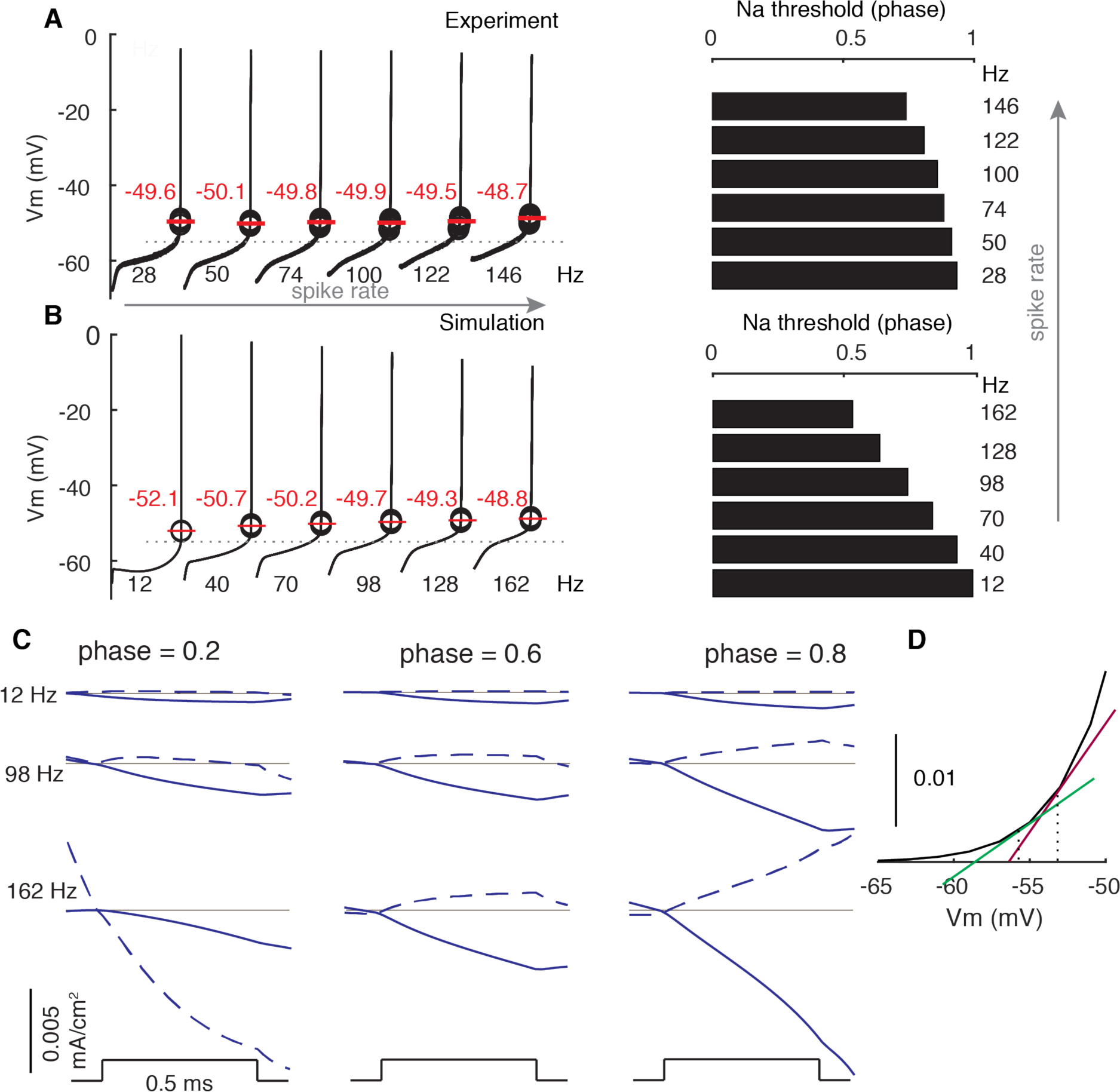
Modulated Subthreshold Membrane Potentials Account for the Rate Adaptation of PRCs. (**A** and **B**) Experimental and simulated voltage trajectories in PCs at different rates. All voltage trajectories are shown from trough to peak within normalized ISIs. The model used (Zang et al., 2018) was not fitted to this specific experimental data. Spike thresholds at different rates are labeled in plots. The Na^+^ activation threshold is defined as -55 mV (stippled line). Right plots show phase dependence of Na^+^-activation threshold on firing rates. (**C**) Stimulus-triggered variations of inward ionic currents (solid) and outward ionic currents (dashed) at different phases and rates. Ionic currents are shifted to 0 (grey line) at the onset of stimulus to compare their relative changes. At phase = 0.2, the outward current is still decreasing due to the inactivation of the large conductance Ca^2+^-activated K^+^ current at 162 Hz. (**D**) Larger slopes of the Na^+^ activation curve at high membrane potentials.

During early phases of all firing rates, membrane potentials are distant from the Na^+^ activation threshold of the transient Na^+^ channel (Fig. 2A, B). The depolarizations to weak stimuli fail to activate sufficient transient Na^+^ channels to speed up voltage trajectories (Fig. 2C). Consequently, phase advances in early phases are small and flat. At later phases, membrane potentials gradually approach and surpass the transient Na^+^ activation threshold. Stimulus-evoked depolarizations activate more Na^+^ channels to speed up trajectories in return. Therefore, phase advances become large and phase-(actually voltage-) dependent. Because high rate-corresponding elevated membrane potentials have larger slopes at the foot of the Na^+^ activation curve, the same ΔV activates more Na^+^ channels and, in addition to the normalization, contributes to larger PRC peaks at high rates (Fig. 2C, D). Under all conditions (except phase = 0.2, 162 Hz), stimulus-evoked depolarizations also increase outward currents, but this increase is smaller than that of inward currents (mainly Na^+^) due to the high activation threshold of K^+^ currents (mainly Kv3) in PCs (Martina et al., 2003; Zang et al., 2018). As the stimulus becomes stronger, it triggers larger depolarizations and the required pre-stimulus membrane potential (phase) to reach Na^+^ activation threshold is lowered. Thus, increasing the stimulus amplitude not only increases PRC peaks, but also shifts the onset-phases of phase-dependent responses to the left (Fig. S1C). In the absence of a dendrite (Fig. S2), the larger amplitude spike is followed by a stronger afterhyperpolarization (Zang et al., 2018) that deactivates K^+^ currents allowing for an earlier depolarization in the ISI, resulting in a completely different PRC.

We further confirmed that the critical role of subthreshold membrane potentials in shaping PRC profiles is not specific to the PC by manipulating PRCs in a modified Traub model (Ermentrout et al., 2001) (Fig. S3 and accompanying text).

### Rate-dependent High-frequency Oscillations

The potential effect of firing rate-caused variations of cellular response properties on population synchrony has been largely ignored in previous studies (Bartos et al., 2002; Brunel and Hakim, 1999; de Solages et al., 2008; Heck et al., 2007; Shin and De Schutter, 2006). Here, we examine whether spike rate correlates with synchrony in the presence of high-frequency oscillations that have been observed in the cerebellar cortex (Cheron et al., 2004; de Solages et al., 2008). We built a biophysically realistic network model composed of 100 PCs with passive dendrites distributed on the parasagittal plane (Witter et al., 2016). Each PC connects to the somas of its 5 nearest neighboring PCs through inhibitory axon collaterals on each side based on experimental data (Bishop and O’Donoghue, 1986; de Solages et al., 2008; Watt et al., 2009; Witter et al., 2016). Rates of each PC are independently driven by parallel fiber synapses, stellate cell synapses, and basket cell synapses (Fig. 3A). More details are in **Methods**.

**Figure 3.**
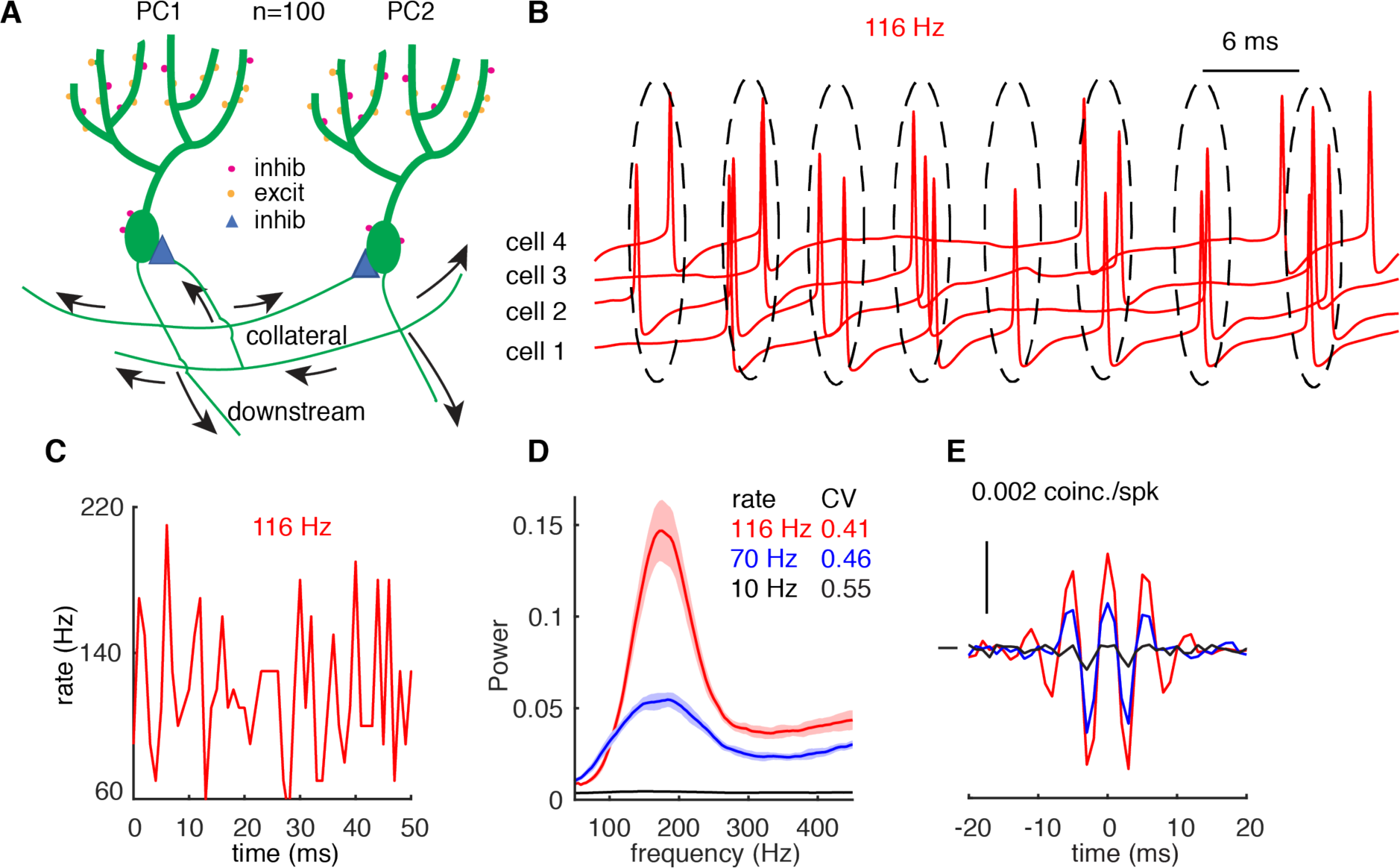
High-frequency Oscillations Show Adaptation to Cellular Firing Rates. (**A**) Schematic representation of the network configuration. (**B**) Example of sampled PC voltage trajectories in the network. (**C**) Example of population rates in the network (time bin 1 ms). (**D**) The power spectrum of population rates of the network at different cellular rates and firing irregularity (CV of ISIs). (**E**) Averaged normalized cross-correlations at different cellular rates.

When the average cellular rate is 116 Hz, PCs in the network tend to fire within interspaced clusters with time intervals of ∼ 6 ms (Fig. 3B). However, individual PCs do not fire within every cluster. Therefore, spikes in the network show intermittent pairwise synchrony on the population level rather than spike-to-spike synchrony (Fig. 3B). Each peak in Fig. 3C corresponds to a ‘cluster’. Based on the power spectrum, the network oscillates at a frequency of ∼ 175 Hz (inverse of the cluster interval, ∼ 6 ms), which is independent of cellular firing rates (116 Hz in red and 70 Hz in blue, Fig. 3D), because oscillation frequency is mainly determined by synaptic properties (Brunel and Hakim, 1999; Brunel and Wang, 2003; de Solages et al., 2008; Maex and De Schutter, 2003). When cellular firing rates increase from 70 Hz to 116 Hz, the power of high-frequency oscillations significantly increases and the peak becomes sharper. When individual PCs fire at low rates (10 Hz), the network fails to generate high-frequency oscillations and each PC fires independently, as evidenced by the flat power spectrum (Fig. 3D). High-frequency oscillations and their firing rate-dependent changes are also reflected in the average normalized cross-correlograms (CCGs) between PC pairs (Fig. 3E). When PCs fire at 70 Hz and 116 Hz, in addition to positive central peaks, two significant side peaks can be observed in the CCGs, suggesting correlated spikes with 0 ms-time lag and ∼ 6 ms-time lag. Amplitudes of the peaks increase with cellular firing rates and disappear when they are low (10 Hz).

In Fig. 3, the variation of cellular rates was driven by synaptic input to demonstrate the rate adaptation of high-frequency oscillations. However, it is difficult to differentiate the relative contribution of PRC shapes and firing irregularity (measured by the CV of ISIs) since they covary with firing rates (Fig. 3D). Therefore, cellular rates were systematically varied by dynamic current injections, which were approximated by the Ornstein–Uhlenbeck (OU) process (Destexhe et al., 2001). This simulation protocol also causes the formation of high-frequency oscillations (Fig. S4).

When PCs fire with low to moderate CV of ISIs, they show loose spike-to-spike synchrony at high rates, and the power peak increases with cellular firing rates. High-frequency oscillations were never observed for low cellular firing rates (Fig. 4A, S4). With high CV of ISIs, spikes are jittered and spike-to-spike synchrony is disrupted (Fig. 4B). Oscillation changes due to firing properties are also reflected in average normalized CCGs. Both central and side peaks increase with the cellular firing rate and decrease with the spiking irregularity. Our results show that small spiking irregularity supports high-frequency oscillations.

**Figure 4.**
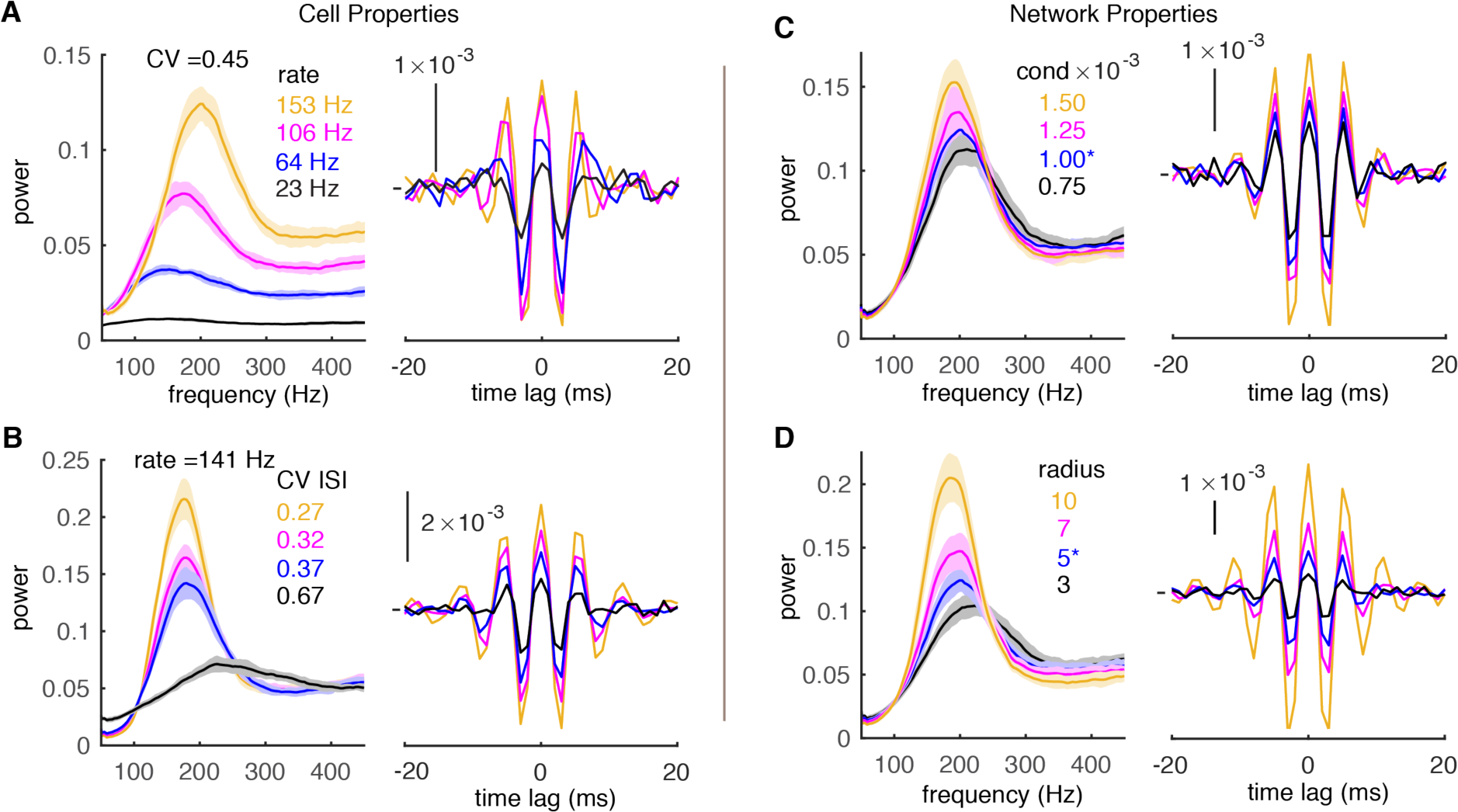
Effect of Cell and Network Properties on High-frequency Oscillations. (**A**) Low cellular firing rates decorrelate the network output in the forms of reduced peaks of power spectrums (left) and CCGs (right). CV ISI is ∼ 0.45. Synaptic conductance is 1 nS and radius is 5. (**B**) Irregular spiking (high CV of ISIs) also decorrelates network. The cellular firing rate is ∼ 141 Hz. Same layout and network properties as in A. (**C**) Small conductance (cond) of inhibitory synapses decorrelates network output. Same layout and network properties as in A with cellular firing rate ∼ 151 Hz and CV ISI ∼ 0.45. (**D**) Short connection radius also decorrelates network output. Same layout and cellular firing properties as C.

At the circuit level, the strength of inhibitory synapses and connection radius are difficult to determine accurately, but their values are critical for the function of axon collaterals. Within the ranges of experimentally reported synaptic conductance and connection radius (de Solages et al., 2008; Fisyunov et al., 2006; Orduz and Llano, 2007; Watt et al., 2009; Witter et al., 2016), the network generates robust high frequency oscillations (Fig. 4C, D). In addition, we find that increasing the conductance of inhibitory synapses or their connection radius increases the power of high-frequency oscillations and make the power spectrum sharper. The increased oscillation power due to connectivity properties is also captured by the larger peaks in CCGs.

Together, our simulation data suggest that the correlation between PC spikes is strong under conditions of low to moderate spiking irregularity, high cellular firing rate, high synaptic conductance, and large connection radius.

### High-frequency Oscillations are Caused by Rate-dependent PRCs

Because both oscillation power and PRC are firing rate dependent, a causal relationship is possible. This is supported by the effect of PRC size on oscillations: decreasing its size leads to weaker oscillations and can even cause weaker oscillations at higher spike rates (Fig. S5). However, it is impossible to manipulate PRC shapes in the complex PC model without greatly affecting other cell and network properties. Therefore, we investigated the effect of rate-dependent PRC shapes in a network of simple coupled oscillators (Kuramoto, 1984; Smeal et al., 2010), where the firing rate specific PRC was used as the coupling term *Z*(*θ*) (see **Methods**). In such a coupled oscillator network, the oscillation power shows a firing rate dependence similar to that of the complex PC network (Fig. 5A, B). This finding demonstrates that the firing rate adaptation of the PRC is sufficient to cause firing rate dependent oscillations.

**Figure 5.**
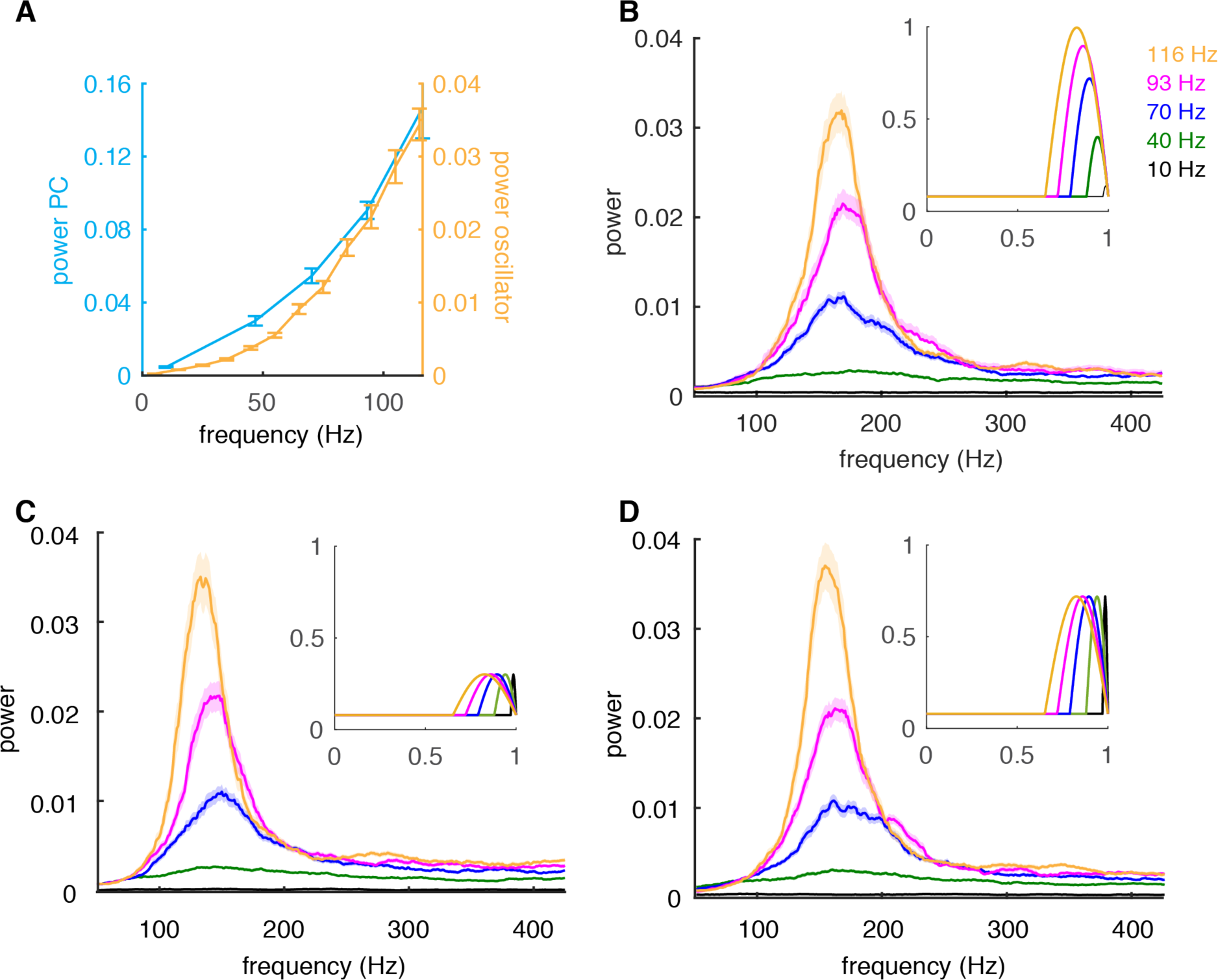
Firing-rate Adaptation of High-frequency Oscillations Is Caused by the PRC. (**A**) Dependence of peak power of high-frequency oscillations in the complex PC network of Fig. 3 (cyan) and in the coupled oscillator network (orange) on cellular firing rate. (**B**) The power spectrum of the coupled oscillator network depends on the cellular firing-rate specific PRC used as coupling term. Inset: firing-rate dependent coupling factors *Z*(*θ*) used (**C**) Same as B but with the peak amplitude of *Z*(*θ*) set to that of the peak of 30 Hz firing rate. (**D**) Same as C for the peak of 70 Hz firing rate.

Next we investigated the specific contribution of the flat part that dominates the PRC at low firing rates versus the late peak with increasing amplitude that appears at higher firing rates. We checked which of these PRC components is responsible for the effect on oscillations by fixing the amplitude of the peak to the value for a specific firing rate. Networks simulated with fixed PRC peak amplitudes show power spectra (Fig. 5C, D) that are very similar to that obtained with the actual PRC (Fig. 5B). An exception is when peak amplitude is very small (for firing frequencies of less than 30 Hz,not shown). The only significant difference between Fig. 5C and 5D is the peak oscillation frequency, which increases with the firing rate for which the amplitude was taken.

In conclusion, the ratio of flat part width to peak width of the firing rate dependent PRC causes the rate dependence of high-frequency oscillations. At low firing rates the dominant flat part suppresses the coupling between oscillators. At high firing rates the coupling increases during the late peak and synchronizes the oscillators, but the strength of oscillation does not depend on peak amplitude in this network.

### Transient Correlations Form Cell Assemblies

Correlation of spiking has often been proposed as a mechanism to form transient cell assemblies (Abeles, 1982; Hebb, 1949; Singer, 1993). This assumes that oscillations can appear and fade rapidly and that they can appear in networks with heterogeneous firing rates. We have previously simulated networks with a range of homogeneous stable cellular rates. Here, we first test whether rate-dependent synchrony still holds when population rates change dynamically. Population rates of the network approximate the half-positive cycle of a 1 Hz sine wave (peak ∼ 140 Hz) with the duration of each trial being 1 sec (Figs. 6A, S6B). We compute shuffle-corrected, normalized joint peristimulus time histograms (JPSTHs) to reflect the dynamic synchrony (Aertsen et al., 1989) (Fig. S6A). The main and the third diagonals of the JPSTH matrix, corresponding to 0 ms-time lag correlation and 6 ms-time lag correlation respectively, are plotted to show the dynamic synchrony at transiently increased rates (bin size is 2 ms, Fig. 6B). At low basal rates, there are no correlations between spikes. Both correlations start to increase ∼ 250 ms after the onset of simulations when the cellular firing rate increases. Closely following rate changes, they decrease again when the cellular rates drop. It demonstrates that axon collateral-caused spike correlations can be achieved transiently to transmit a correlation code conjunctive with temporal cellular firing rate increases. Similar results were obtained for a faster change of population rates (2.5 Hz sine wave, Fig. S7).

**Figure 6.**
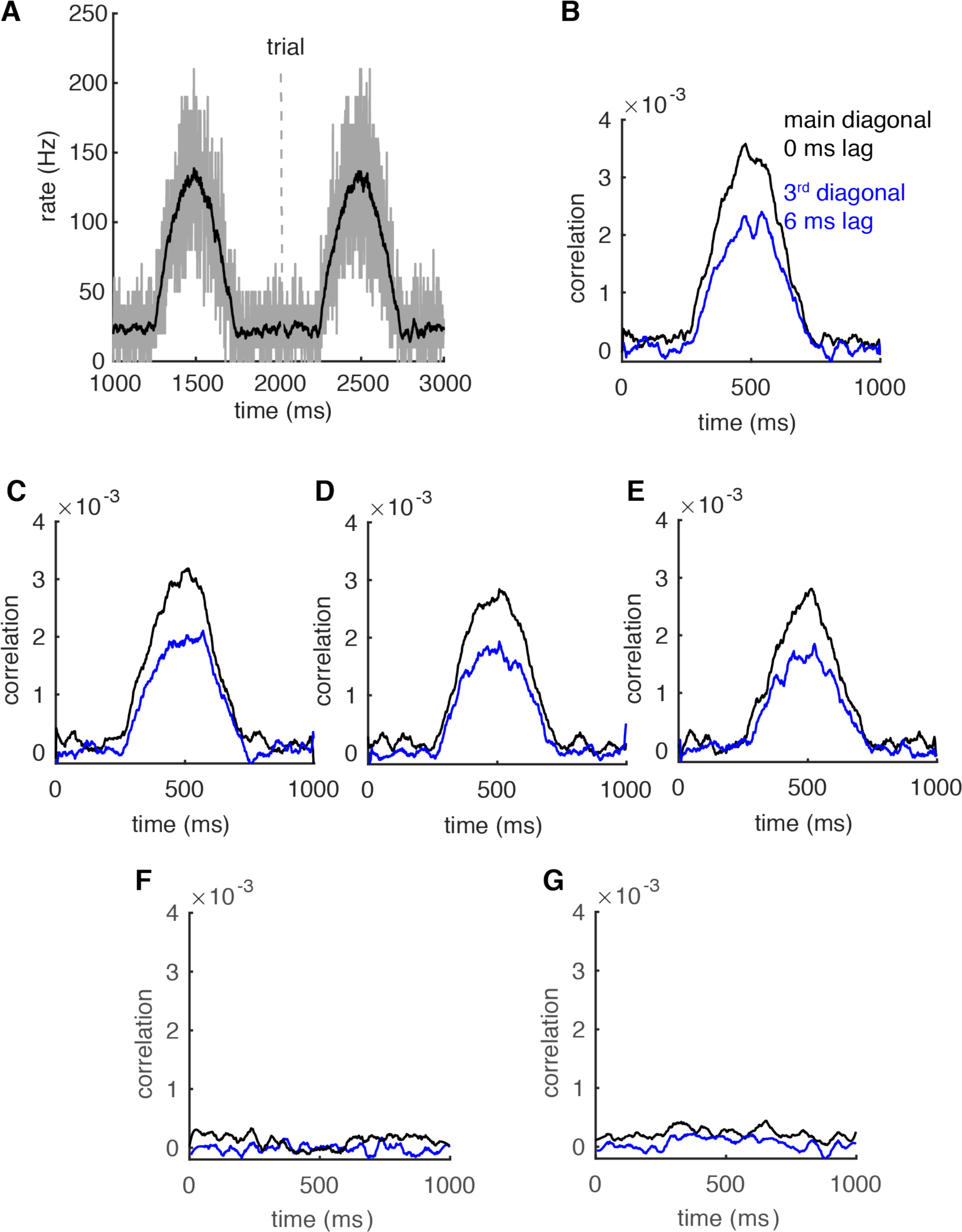
Correlations Can be Transient and Robust to Heterogeneous Spike Rates. (**A)** Population spike rates of PCs. (**B)** The 0 ms- and 6 ms-time lag correlations increase with population rates. (**C-E**) The rate-dependent correlation is robust to heterogeneous cellular rate changes. From **C** to **E**, the number of decreased rate cells increases from 10 to 30. (**F**) Correlations between decreased-rate neurons in the network (n = 30). (**G**) Correlations between increased-rate neurons and decreased-rate neurons (n = 30 for each group).

Although it remains unclear whether the population of PCs converging onto a same cerebellar nuclei (CN) neuron are homogeneous or heterogeneous (Uusisaari and De Schutter, 2011), simultaneous bidirectional PC rate changes have been observed during cerebellum-related behaviors (Chen et al., 2016; Herzfeld et al., 2015). It is very likely that neighboring PCs show heterogeneous spike rate changes (Hong et al., 2016), which can reduce spike correlations (Markowitz et al., 2008). Therefore, we distributed 10-30 extra cells with decreasing spike rates (Fig. S6B) in the network to test the effect of heterogeneous neighboring rate changes on transient correlations. They were randomly scattered among the cells with increasing rates. Spike correlations still become larger for the subgroup of PCs showing increased cellular rates, despite a slight decrease of the correlation amplitude when more cells decrease their spike rates (Figs. 6C-E). Moreover, the spiking in PCs with decreased firing rates is not correlated (Fig. 6F), nor is it correlated with oscillating increased-rate PCs (Fig 6G), making the assembly formation specific to fast spiking PCs. The results suggest that a population of PCs with increased spike rates can form a correlated assembly that will strongly affect downstream neurons even when it is surrounded by non-correlated neighboring PCs with decreased spike rates.

## Discussion

In this work, we reproduced the firing rate-dependent PRC of PCs and dissected the underlying biophysical mechanisms. Next we explored the role of these PRCs in synchronizing spikes in cerebellar PCs and how they can support the formation of transient assemblies.

### Biophysical Mechanisms Underlying Rate-dependent PRCs

The profiles of neuronal PRCs are rate dependent (Couto et al., 2015; Ermentrout et al., 2001; Gutkin et al., 2005; Phoka et al., 2010; Tsubo et al., 2007). Cerebellar PCs exhibit a transition from small, phase-independent responses to large, phase-dependent type-I responses with increasing rates (Couto et al., 2015; Phoka et al., 2010), but the mechanism was unknown (Akemann and Knopfel, 2006; Couto et al., 2015; De Schutter and Bower, 1994; Khaliq et al., 2003; Phoka et al., 2010). This work reproduces and explains the experimentally observed rate adaptation of PRCs. Note that the slight increase of PRCs in the very narrow late phase in our model (low rate, Fig. 1B) may be annihilated by noise in spontaneously firing neurons (Couto et al., 2015; Phoka et al., 2010).

Compared with previous work emphasizing the slow deactivation of K^+^ currents in cortical neurons (Ermentrout et al., 2001; Gutkin et al., 2005), here we demonstrate the role of rate-dependent subthreshold membrane potentials and their corresponding activation of Na^+^ currents. In both pyramidal neurons and PCs, spike rates cause significant variation of the subthreshold membrane potential during the ISI (Rancz and Hausser, 2010; Tsubo et al., 2007). In response to a stimulus, both Na^+^ and K^+^ currents are activated. In PCs, the main K^+^ current is high-threshold activated (Martina et al., 2003; Zang et al., 2018); therefore, depolarization-facilitated Na^+^ currents dominate, causing larger normalized PRCs at high rates (Fig. 2). However, previous PC models (Akemann and Knopfel, 2006; Couto et al., 2015; De Schutter and Bower, 1994; Khaliq et al., 2003; Phoka et al., 2010) included low-threshold-activated K^+^ currents, which counteract facilitated Na^+^ currents. In the original Traub model, slow deactivation of K^+^ currents and consequent hyperpolarization synergistically reduce the normalized PRC peaks at high rates (Ermentrout et al., 2001; Gutkin et al., 2005). By minimally modifying the Traub model, elevated subthreshold membrane potentials generate larger normalized PRC peaks at high rates (Fig. S3).

### The Evidence Supporting Rate-dependent Correlations

Rate-dependent synchrony in the cerebellum has been demonstrated for Golgi cells (van Welie et al., 2016) but not, as yet, for PCs. However, careful analysis of previous experimental data in the cerebellum provides some evidence to support our findings. In the work of de Solages et al. (2008), units with lower average rates (<10 Hz) did not exhibit significant correlations between neighboring PCs, for unknown reasons. This can be explained by the small flat PRCs at low rates. Under extreme conditions, when the PRC is constantly 0 (equivalent to disconnection), no correlations can be achieved (Figs. 3-6). Additionally, the experimental oscillation power increased by the application of WIN 55,212-2, which was intended to suppress background excitatory and inhibitory synapses (de Solages et al., 2008). The increased power could be due to more regular spiking after inhibiting the activity of background synapses (Fig. 4B). However, it could also be caused by increased spike rates (Fig. 4A), because this agent also blocks P/Q type Ca^2+^ channels and consequently P/Q type Ca^2+^-activated K^+^ currents, to increase spike rates (Fisyunov et al., 2006). Similarly, enhanced oscillations have also been observed in calcium-binding protein gene KO mice, which have significantly higher simple spike rates (Cheron et al., 2004). A more systematic experimental study of the firing rate dependent appearance of loose simple spike synchrony among PCs and its relation to behavior would be required to confirm these predictions.

The rate-dependent correlations observed in this study are different from those reported previously by de la Rocha et al. (2007). In that study, common input mediated correlation increased rapidly with increasing rate at low firing rates in pyramidal cells (their Fig. 1e) and in integrate-and-fire models (their Fig 2c), while the PRC mediated correlations in our study for inhibitory coupling only appear at much higher firing rates (Fig. 5A).

### Down-stream effects of PC assemblies

PCs inhibit their target neurons in the CN, which in turn form the only cerebellar output. It is difficult to finely regulate CN firing rates with inhibition only, because it operates over the narrow voltage range between resting potentials and GABA_A_ reversal potentials. Two solutions for this problem have been proposed. The first is that synchronized pauses of PC firing will release CN neurons from inhibition, leading to rebound firing (De Schutter and Steuber, 2009; Lee et al., 2015). There is strong evidence that this mechanism works in controlling the onset of movement in the conditioned eyeblink reflex (Heiney et al., 2014) and in saccade initiation (Hong et al., 2016). The other solution provides a more continuous rate modulated CN output by time-locking of CN spikes to PC input. Several experimental studies have demonstrated that partial synchronization of afferent PC spiking can time-lock the spikes of CN neurons to their input (Gauck and Jaeger, 2000; Person and Raman, 2012). The ability to rapidly increase the correlation level within a subgroup of PCs with increased firing rates (Fig. 6) is therefore predicted to have a strong effect on CN spiking. Moreover, this does not require strong synchronization. Similar results were observed when jitter higher than the few ms predicted by our network model (Fig. 3B) was applied to the synchronous PC input (Gauck and Jaeger, 2000).

### Advantages of Transient PC assemblies

The actual convergence and divergence of PC axons onto CN neurons remains a controversial topic in the literature. There are roughly ten times more PCs than CN neurons and PC axons branch extensively leading to computed convergence values ranging from 20 to over 800 (Uusisaari and De Schutter, 2011), though many authors have recently converged on the compromise of ∼50 (Person and Raman, 2012). If CN neurons just average the activity of all afferent PCs, much of the potential information generated by the large neural expansion in cerebellar cortex would be lost. Our PC network with parameters that fall within physiological ranges can rapidly generate and disrupt oscillations based on the cellular firing rates (Figs. 3,6), with no need of increasing afferent input correlation. Note that rate-related synchrony can also be achieved via common synaptic inputs (Heck et al., 2007), gap junctions (Middleton et al., 2008), and ephaptic coupling (Han et al., 2018), when connections are weak. This means that transiently correlated PC assemblies can form and disappear quickly. Such assemblies, even if consisting of only a few PCs (Person and Raman, 2012), can finely control spiking in CNs. Because the assemblies can consist of variable subsets of afferent PCs to a CN neuron, this greatly expands the information processing capacity of the cerebellum.

## Conclusion

We have shown that firing-rate dependent PRCs can cause firing-rate dependent oscillations at the network level. Such a mechanism supports the rapid formation of transient neural assemblies in cerebellar cortex.

## Acknowledgement

The authors thank for helpful suggestions from Drs. Eve Marder, Tomoki Fukai and Sergio Verduzco to improve the manuscript and for the language editing by Steven Douglas Aird.

## Methods

The detailed PC model and the interconnected network model were implemented in NEURON 7.5 (Carnevale and Hines, 2006). The Traub model was implemented in MATLAB. The code used in this work will be available from ModelDB.

### PRC Computations

Our recently developed compartment-based PC model was used (Zang et al., 2018). To compute the PRCs in Fig. 1, brief current pulses with a duration of 0.5 ms and an amplitude of 50 pA were administered at the soma at different phases of interspike intervals. The resulting perturbed periods were then used to calculate phase advances by (Ermentrout et al., 2001):

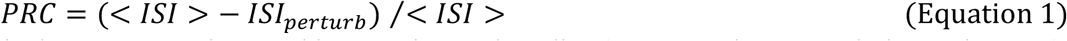

This is the same equation used in experimental studies (Couto et al., 2015; Phoka et al., 2010), to facilitate comparison. Different cellular rates were achieved by somatic holding currents (Couto et al., 2015; Phoka et al., 2010). To compute PRCs in response to negative stimuli, the amplitudes of the pulses were changed to - 50 pA. To compute PRCs of our PC model with passive dendrites, only H current and leak current were distributed on the dendrites with the same parameters as in the active model (Zang et al., 2018). The Traub model (Traub et al., 1999) was implemented according to the work of Ermentrout et al (Ermentrout et al., 2001; Gutkin et al., 2005). In the modified version of this model, the conductance of the kdr current was reduced from 80 to 40. Activation and deactivation rates of this current were shifted to the right by 30 mV, α_n_(v) = 0.032*(v+22)/(1-exp(-(v+22)/5)); β_n_(v) = 0.5*exp(-(v+27)/40); the conductance of AHP current was increased from 0 to 0.1.

### Network Simulations

We implemented our recurrent inhibitory PC layer network using the Watts-Strogatz model (Watts and Strogatz, 1998) to avoid boundary effects. To reduce simulation time, we used the PC model with passive dendrites, which exhibits similar rate-dependent PRCs as the PC model with active dendrites (Fig. S1B). In the baseline version of the network, 100 PCs were distributed on the parasagittal plane (Witter et al., 2016), corresponding to 2 mm of folium with a distance of 20 µm between neighboring PC soma centers. 100 PCs are within the estimated range of PCs converging to a same cerebellar nuclei neuron (Person and Raman, 2012). Each PC was connected to its nearest 2*radius neighboring PC somas and connections had 0 rewiring probability. The PCs were interconnected, according to anatomical data showing collaterals present toward both the apex and the base of the lobule with only slight directional biases (Witter et al., 2016). The baseline value of radius was 5 within the range of experimental estimates (Bishop and O’Donoghue, 1986; de Solages et al., 2008; Watt et al., 2009; Witter et al., 2016). The inhibitory postsynaptic current (IPSC) was implemented using the NEURON built-in point process, Exp2Syn. G = weight * (exp(-t/τ_2_) - exp(-t/τ_1_)), with τ_1_ = 0.5 ms (rise time) and τ_2_ = 3 ms (decay time). The reversal potential of the IPSC was set at -85 mV (Watt et al., 2009). The conductance was 1 nS (de Solages et al., 2008; Orduz and Llano, 2007; Witter et al., 2016). The delay between onset of an IPSC and its presynaptic spike timing was 1.5 ms (de Solages et al., 2008; Orduz and Llano, 2007; Witter et al., 2016). To test the effect of rate-dependent PRCs on high-frequency oscillations, we varied the cellular rates in two paradigms. In the first paradigm (Fig. 3), each PC is contacted by 4,000 excitatory parallel fiber synapses (PF, on spiny dendrites), 18 inhibitory stellate cells (STs, on spiny dendrites) and 4 inhibitory basket cells (BSs, on the soma). Activation of excitatory and inhibitory synapses in each PC was approximated as an independent Poisson process with different rates. We simulated 5 conditions: PC rate = 10 Hz when PF rate = 0.27 Hz, ST rate = 14.4 Hz, BS rate = 14.4 Hz; PC rate = 47 Hz when PF rate = 1.62 Hz, ST rate = 28.8 Hz, BS rate = 28.8 Hz (used in Fig. 5); PC rate = 70 Hz when PF rate = 2.16 Hz, ST rate = 28.8 Hz, BS rate = 28.8 Hz; PC rate = 93 Hz when PF rate = 2.7 Hz, ST rate = 28.8 Hz, BS rate = 28.8 Hz (used in Fig. 5); PC rate = 116 Hz when PF rate = 3.24 Hz, ST rate = 28.8 Hz, BS rate = 28.8 Hz.

To more systematically explore different factors regulating network outputs we used a second paradigm (Figs. 4, S4, S5). Cellular rates of each PC were manipulated by injecting stochastic currents on the soma. The stochastic current was approximated by the commonly used Ornstein-Uhlenbeck random process (Destexhe et al., 2001), 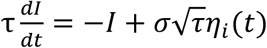. σ represents the amplitude of the fluctuation; *η*_*i*_ represents uncorrelated white noise with unit variance; τ = 5 ms. In this paradigm, we systematically varied the rates and firing irregularities of PCs (CV of ISIs) to explore their importance for network output. Due to the intrinsic relationship between CV of ISIs and firing rates, a larger σ is required for large *I* to get a same CV of ISI. Phase response is a result of input current and response gain of the cell. We reduce the phase response by halving the input current (synaptic conductance) to achieve a smaller response at high firing rates (Fig. S5). The conductance of inhibitory synapses was tested with the values of 0.75, 1.0, 1.25 and 1.5 ns in Fig. 4C. We also explored the effect of connection radius with the values of 3, 5, 7 and10 in Fig. 4D.

To test a spatio-temporally increased correlation, we randomly distributed extra 10-30 PCs with decreased cellular rates into the original network (Figs. 6, S7), including 100 increased-rate cells. Their mean population firing rates are shown in Fig. S6B.

### Coupled Oscillator Model

The model comprises 100 neurons that are randomly connected to each other with connection probability of *p* = 0.75 (Fig. 5). The “subthreshold dynamics” of individual neurons is given by the phase equation

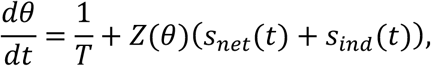

where *θ* is a phase variable ranging from 0 to 1. *T* is a period of the oscillation. *Z*(*θ*) is a PRC. *s*_*ind*_ and *s*_*net*_ are the individual and network input, respectively. At *θ*=1, the model cell “spiked.” Then, *θ* was reset to *θ*-1 and the spike was added to the spike train variable (see below).

*Z*(*θ*) is given by

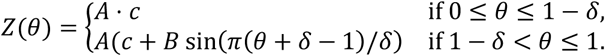

Here *c* represents the flat part of the Purkinje cell PRC and the other term represents a “bump” around *θ* = 1. We found that the bump width is ∼3 ms in time regardless of the firing rate, and set *δ* = 3 ms/*T*. We also used *c* = 0.08 and *A* = 12.

In the case when the model PRC scales as the PC PRC (Fig. 5B), *B* = *f*_*amp*_(1/*T*) where *f*_*amp*_(*r*) is a normalized PRC amplitude given a baseline firing rate *r* in Fig. 1C. In Fig 4C and D with no amplitude scaling of the PRC, *B* = *f*_*amp*_(30 Hz) and *B* = *f*_*amp*_(70 Hz) are used, regardless of *T*, respectively.

*s*_*ind*_(*t*) is given by the Ornstein-Uhlenbeck (OU) process, *∂*_*t*_*s*_*ind*_ = −*s*_*ind*_/*τ* + σ_0_*ζ*, where *ζ* is a Wiener process based on the standard normal distribution. We used *τ* = 3 ms and σ_0_ = 0.2.

*s*_*net*_(*t*) is given by

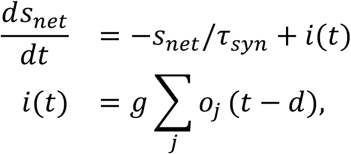

where *j* represents other neurons connected to each cell, and *o*_*j*_(*t*) is a spike train of the cell *j. d* = 1.5 ms is a synaptic delay. *g* = 20 is a connection weight, and *τ*_*syn*_= 3 ms is a decay time for the synaptic current.

We used the forward Euler method with a time step of 0.025 ms to integrate the subthreshold equation, while we also confirmed that our results did not change if we use 0.0125 ms. The OU processes were integrated with the same time step and backward Euler method.

### Data Analysis

The power spectrum of the spike trains of the network was estimated by Welch’s method, which calculates the average of the spectra of windowed segments (window size 128 points). In each trial under each specific stimulus condition, the length of the signal was 2 sec, with a time resolution of 1 msec. The final result was the average of 14 trials.

To compute the cross-correlogram (CCG) under each specific stimulus condition, we first computed pairwise correlations between the spike trains of two neurons and then corrected them by shift predictors, which removed the ‘chance correlations’ due to rate changes. Then correlations were divided by the triangular function Θ(*τ*) = *T* − |*τ*| and 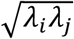. T was the duration of each trial and *τ* was the time lag. Θ(*τ*) corrects for the degree of overlap between two spike trains for each time lag *τ. λ*_*i*_ was the mean firing rate of neuron *i* (Kohn and Smith, 2005). Finally, the CCGs between all pairs in the network were averaged to reflect the population level spike correlations. Thus, similar with previous work (Heck et al., 2007), the computed CCGs reflect the ‘excess’ correlation caused by axon collaterals in our work.

To measure the dynamic correlation over the time course of the stimulus, we computed JPSTHs (Aertsen et al., 1989). We first picked two neurons from our network and aligned their spike-count PSTHs to stimulation onset with 2-ms time bins in each trial (larger time bins annihilated the positive peaks due to the significant negative correlations in paired spikes, see CCGs in Figs. 3,4). We constructed the JPSTH matrix by taking each stimulus trial segment and plotting the spike counts of one cell on the horizontal and one on the vertical. If there is a spike from neuron i at time *x*, and a spike from neuron j at time *y*, one count will be added to the matrix index (x,y). By repeating this process for different trials, we got a raw matrix for a cell pair i and j. Then by the shift-predictor (repeated previous steps with shuffled stimulation trails), we removed correlations due to co-stimulation caused firing rate changes. Next step, we normalized the JPSTH by dividing with the product of standard deviations of the PSTHs of each neuron. To measure the correlation of the assembly, we averaged JPSTH between all non-repeated cell pairs in the defined “assembly” of our network (Oemisch et al., 2015). The corrected matrix values become correlation coefficients, with values between -1 and +1. The main diagonal of the JPSTH matrix provides a measure of time-varying 0-ms time lag correlations and the third main diagonal (2-ms time bin) provides a measure of 6-ms time lag correlations. Due to the small-time bin we used, we simulated 1992 trials (for Fig. 6 B-E) to compute JPSTH between PC pairs and smoothed the JPSTHs for visualization purpose. Due to the small number of decreased-rate neurons in the network, we simulated 26112 trials to compute Fig. 6F, G (30 decreased-rate neurons). When decreased-rate neuron numbers are 10 and 20 (Fig. 6C, D), we did not compute their correlations due to the computational challenge. For Fig. 6G, we randomly picked 30 from 100 increased-rate neurons to make pairs with 30 decreased-rate neurons.

## Supplementary material and figures

**Figure S1.**
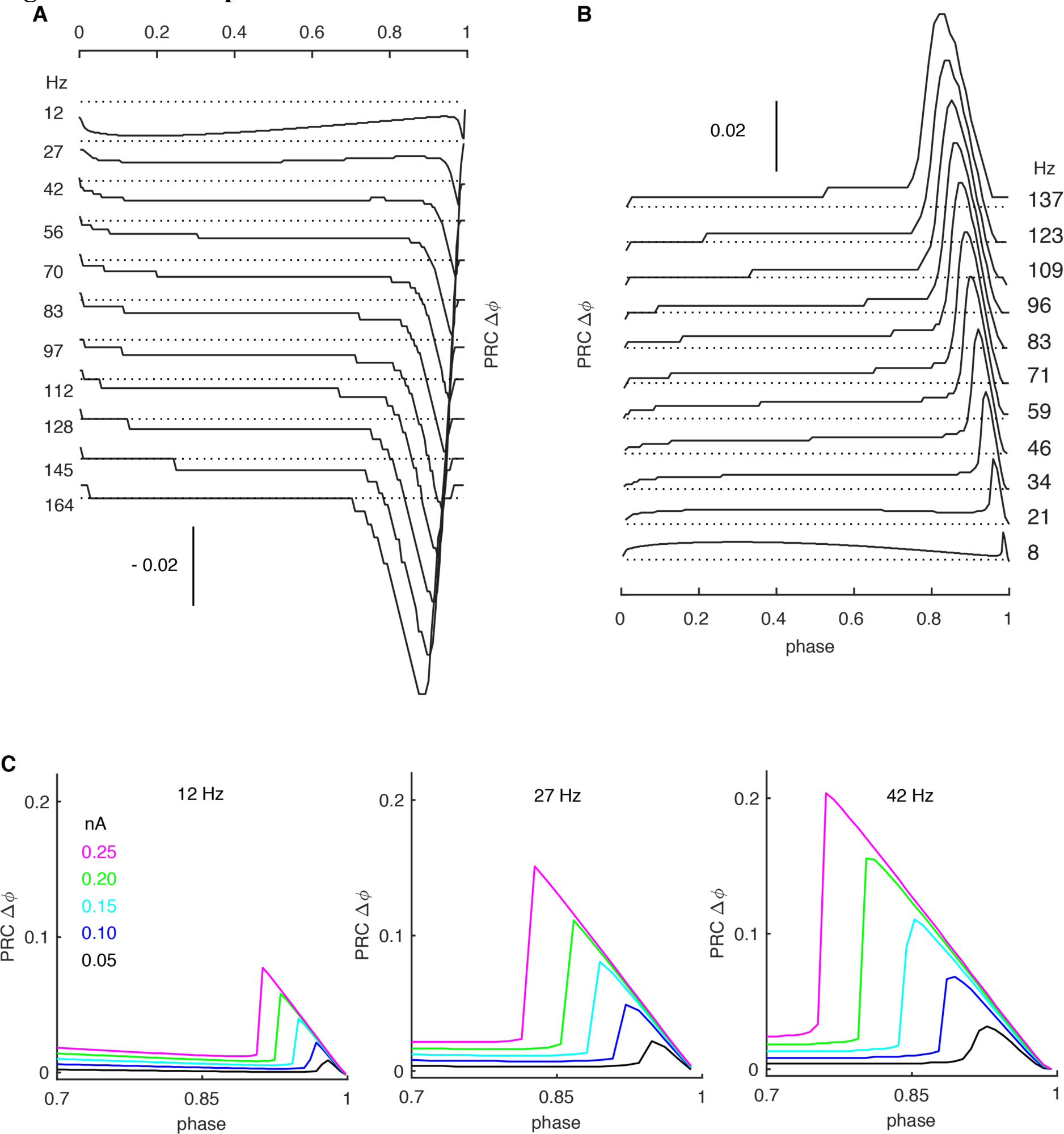
Rate-dependent PRCs. (**A**) Negative stimulus-triggered responses (phase delay) parallel positive stimuli (see Fig .1). Onset-phases of phase-dependent responses shift left at high rates with gradually larger amplitudes. (**B**) The PC model with passive dendrites shows similar rate adaptation as in PC models with active dendrites (Fig. 1). (**C**) Larger stimulus amplitudes increase the peak of the phase-dependent PRCs and shift their onset phases to the left. Simulation results at rates of 12-, 27- and 42-Hz are shown with increased stimulus amplitudes from 0.05 nA to 0.25 nA.

**Figure S2.**
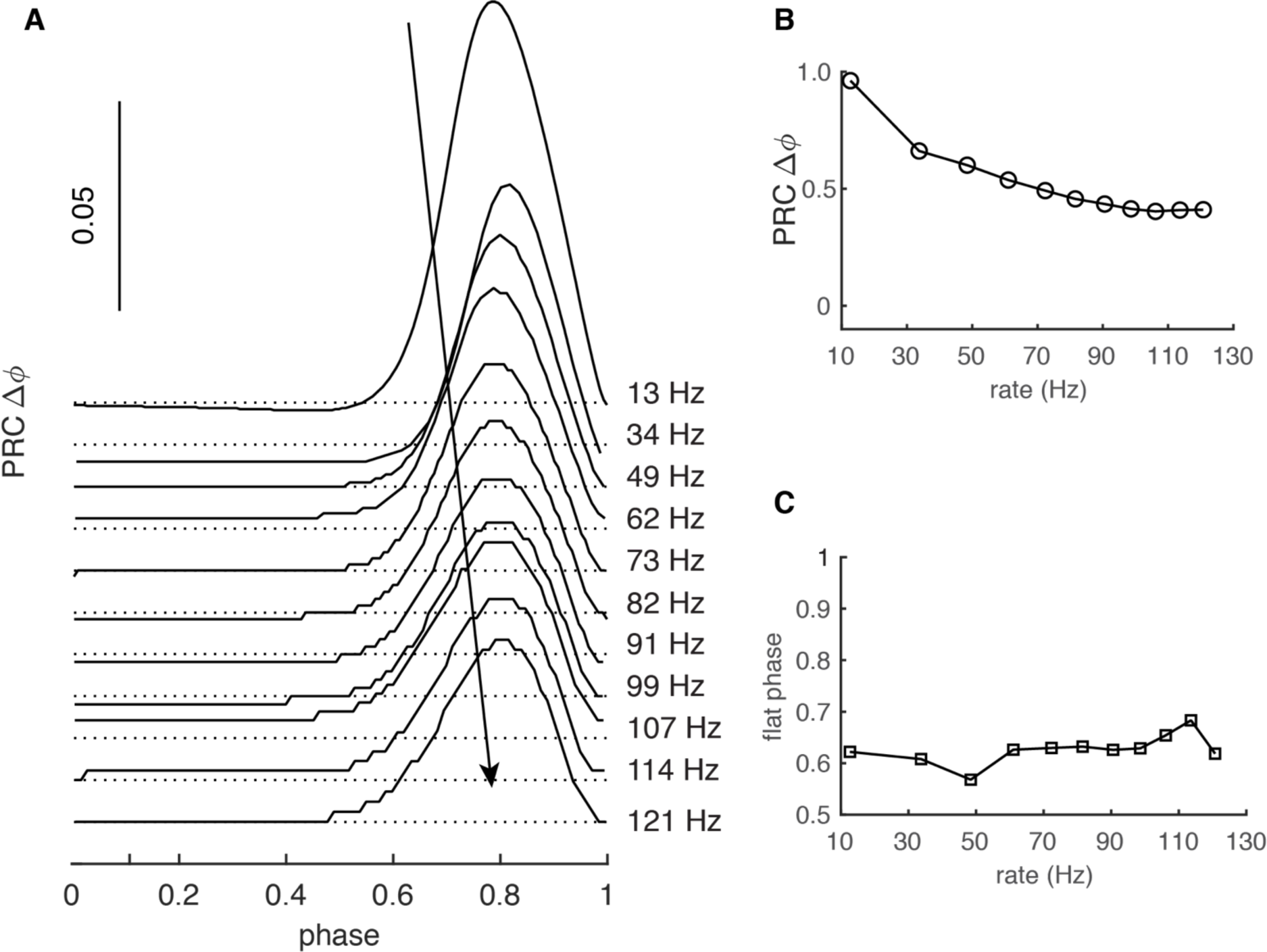
Rate-dependent PRCs are influenced by the dendrite. PRC obtained in the PC model without its dendrite. (**A**) The PRC has a different rate adaptation: the peak has a constant width and its amplitude decreases with firing rate. (**B)** Normalized peak amplitudes at different firing rates. (**C**) The flat phase has a constant duration.

**Figure S3.**
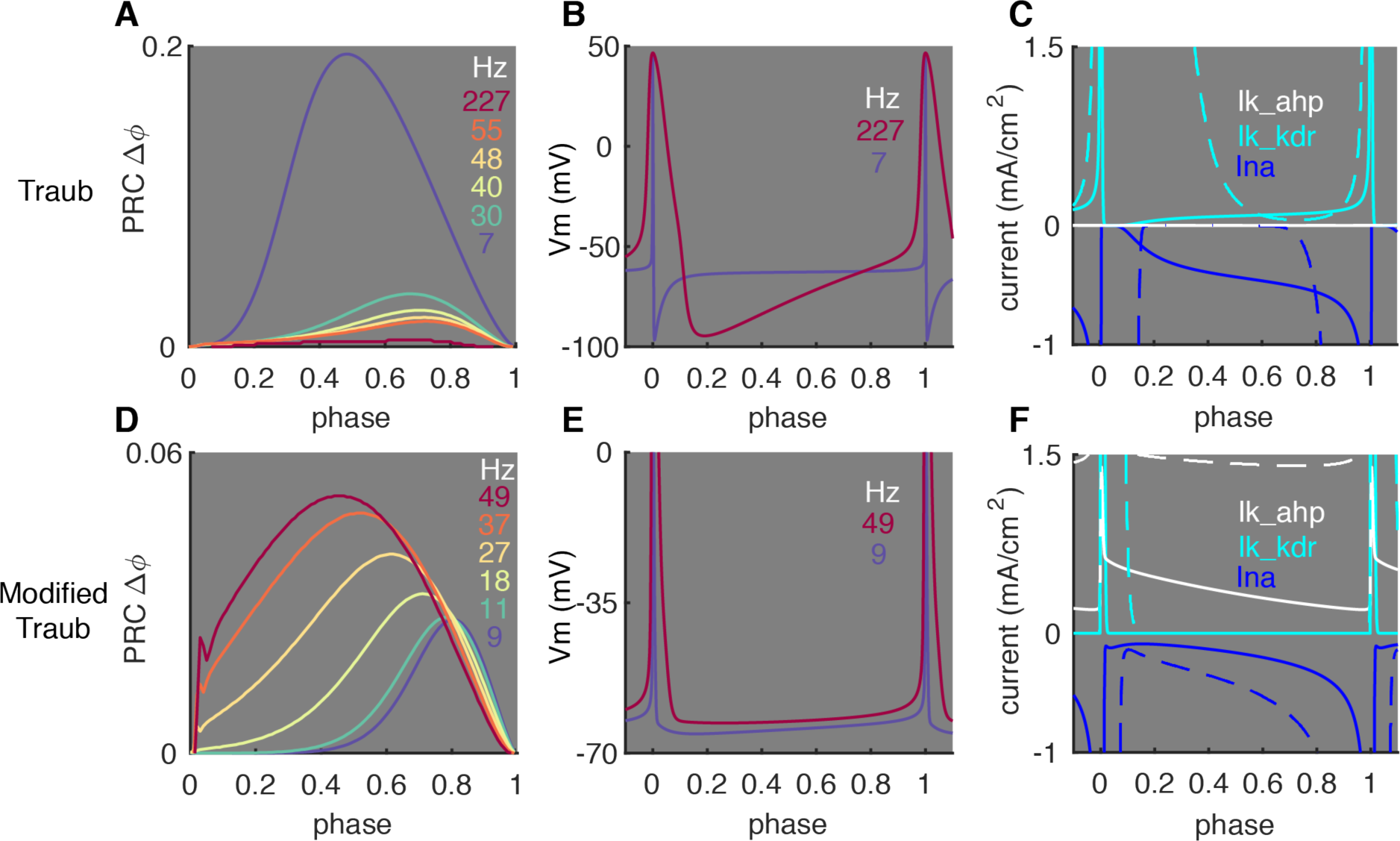
Effect of Subthreshold Membrane Potentials on Shaping PRCs. We examined whether the critical role of subthreshold membrane potentials in shaping PRC profiles (Fig. 2) also applies to other neuron types. A frequently used pyramidal neuron model, the Traub model (Ermentrout et al., 2001) was tested. It shows an opposite rate adaptation of PRCs compared to PCs (Fig. S3A). In the Traub model, responses become smaller and relatively phase-independent at high firing rates. This demonstrates that the normalization used in Eq. 1 does not always lead to increasing PRC peak amplitudes for smaller ISIs. These PRC shapes can be explained by significantly lower subthreshold membrane potentials at high rates, compared to PCs, (Fig. S3B). This is due to the accumulation of delayed rectifier K^+^ current (kdr, Fig. S3C), which has a low activation threshold and large conductance. The lower subthreshold membrane potentials are far below the Na^+^-activation threshold, making responses to weak stimuli passive at high firing rates. Accordingly, PRCs in the model become smaller and relatively phase-independent at high rates. We minimally modified the Traub model by reducing the conductance of the kdr current, raising its activation threshold and increasing the AHP current (details in **Methods**) (Fig. S3D-F). With these modifications, subthreshold membrane potentials are significantly elevated at high firing rates lower than 110 Hz. Accordingly, onset-phases of phase-dependent responses shift left and peaks increase at high rates. These simulation results show that spike rate-dependent subthreshold membrane potentials and their effect on nonlinear activation of Na^+^ currents can be crucial in shaping neuronal PRC profiles in many types of neurons. Surprisingly, when firing rates are higher than 110 Hz, PRCs decrease again, suggesting other undetermined biological mechanisms may be involved in determining the phase responses. (**A**) Rate adaptation of PRCs in the original Traub model. (**B**) Lowered ISI membrane potential at high rates. (**C**) Comparison of ionic currents at low (solid, 7 Hz) and high (dashed, 227 Hz) rates. (**D**) Rate adaptation of PRCs in the modified Traub model. (**E**) Elevated ISI membrane potential at high rates. (**F**) Comparison of ionic currents at low (solid, 9 Hz) and high (dashed, 49 Hz) rates. In **C** and **F**, current peaks are truncated to show currents during ISIs. In **E**, spike peaks are truncated to show the elevated ISI membrane potential at high rates.

**Figure S4.**
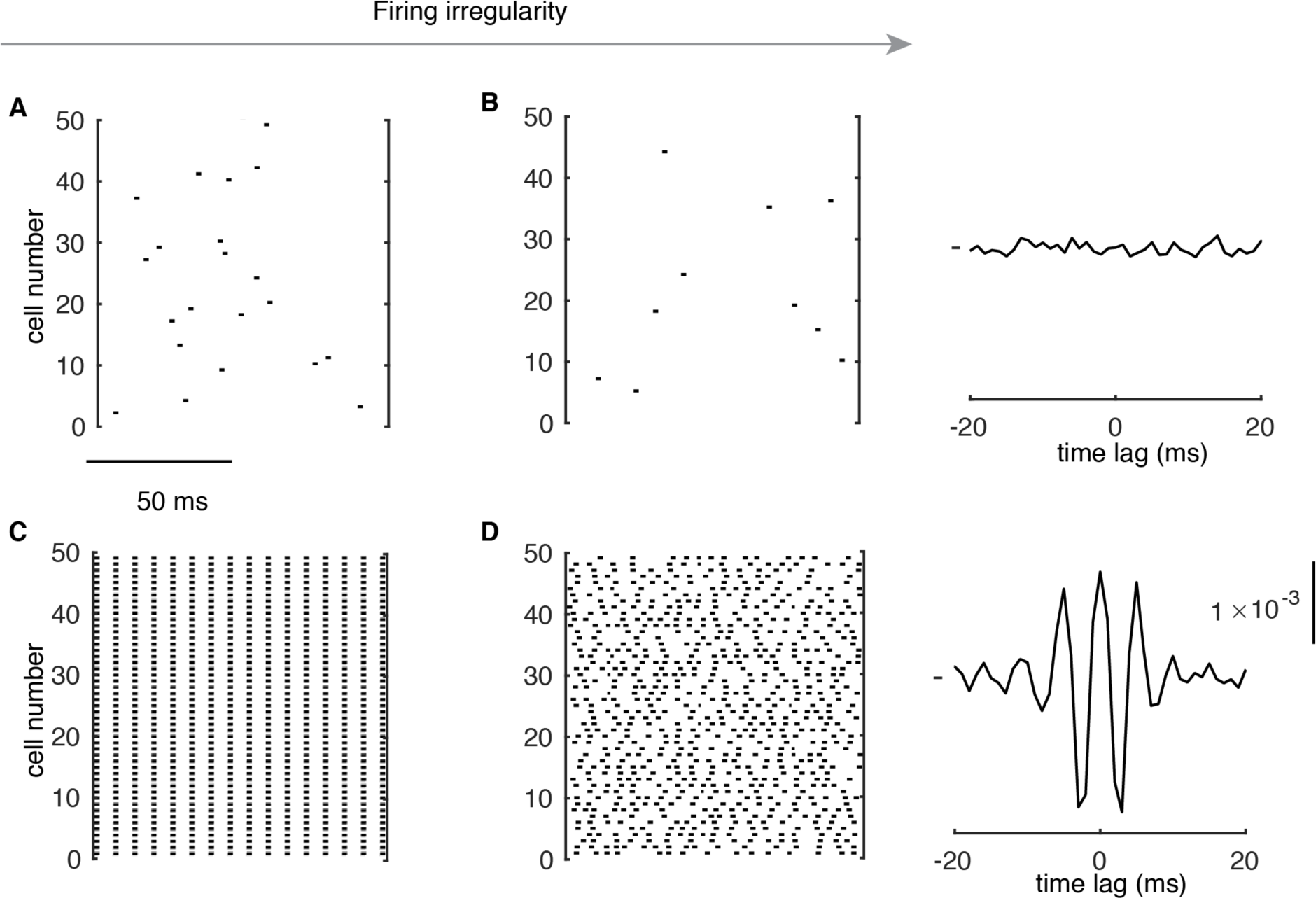
Formation of High-frequency Oscillations at High Rates. (**A**&**B**) Raster plots of random PC spikes when they fire regularly (CV ISI ∼ 0.07) and irregularly (CV ISI ∼ 0.4) at low rates (here, ∼ 2 Hz in the network, but 12 Hz in isolated cells). In the right plot of **B**, average CCG is shown. (**C**) PCs show spike-to-spike synchrony when they fire regularly (CV ISI ∼ 0.02) at high rates (154 Hz). (**D**) PC spikes show high-frequency oscillations when they fire irregularly (CV ISI ∼ 0.44) at high rates (153 Hz). The right plot of **D** shows the average CCG with a significant central peak and side peaks.

**Figure S5.**
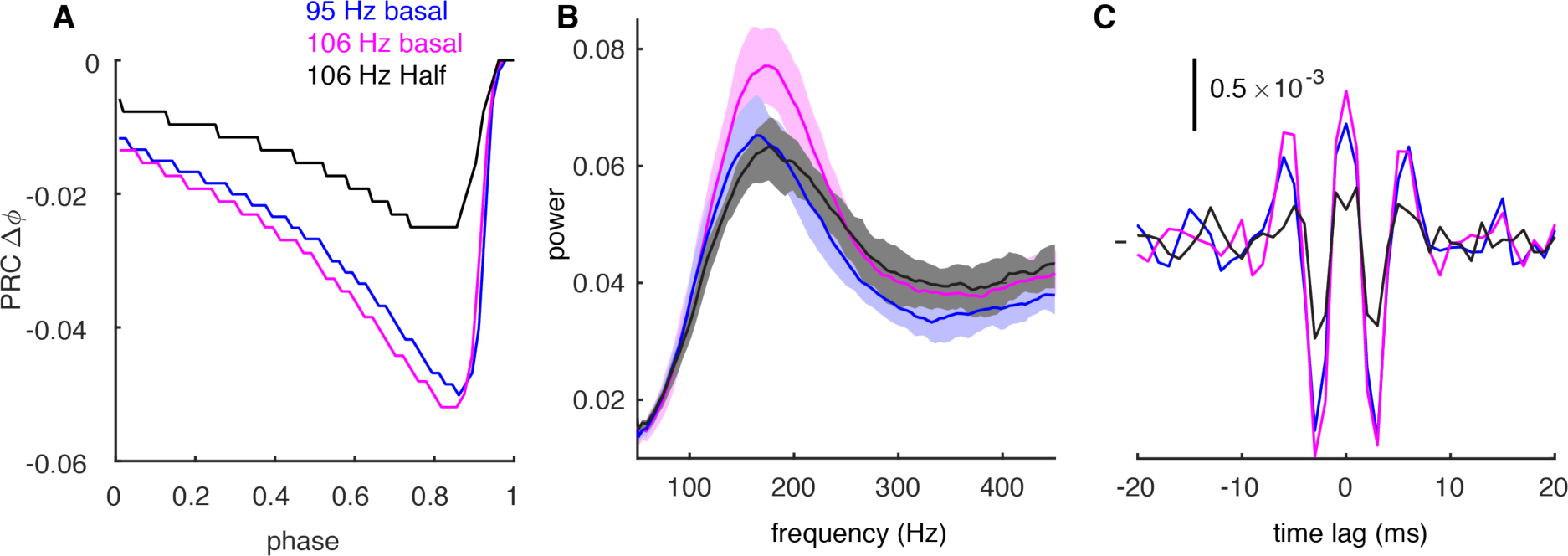
Decreased PRC at High Firing Rates Can Weaken Oscillations. **A**. PRCs for negative synaptic stimulus when PCs fire at 95 Hz (blue) and 106 Hz (purple) driven by OU process with basal values of synaptic conductance. Reduced PRC at 106 Hz was achieved by 50% of basal synaptic conductance (black). **B**. The power spectrum of spike trains with the cellular rate of 95 Hz (blue, basal conductance), 106 Hz (purple, basal conductance) and 113 Hz (black, 50% of basal conductance, firing rate increased to 113 Hz due to the reduced inhibition, with other conditions the same as 106 Hz with basal conductance). In all cases, the CV of ISI is 0.45. The power spectrum at high firing rates gets flatter with lower amplitude when the PRC amplitude is reduced (black trace). **C**. The CCGs of spike trains with the same condition as in B. Central and side peaks reduce at high firing rates when the phase response is smaller.

**Figure S6.**
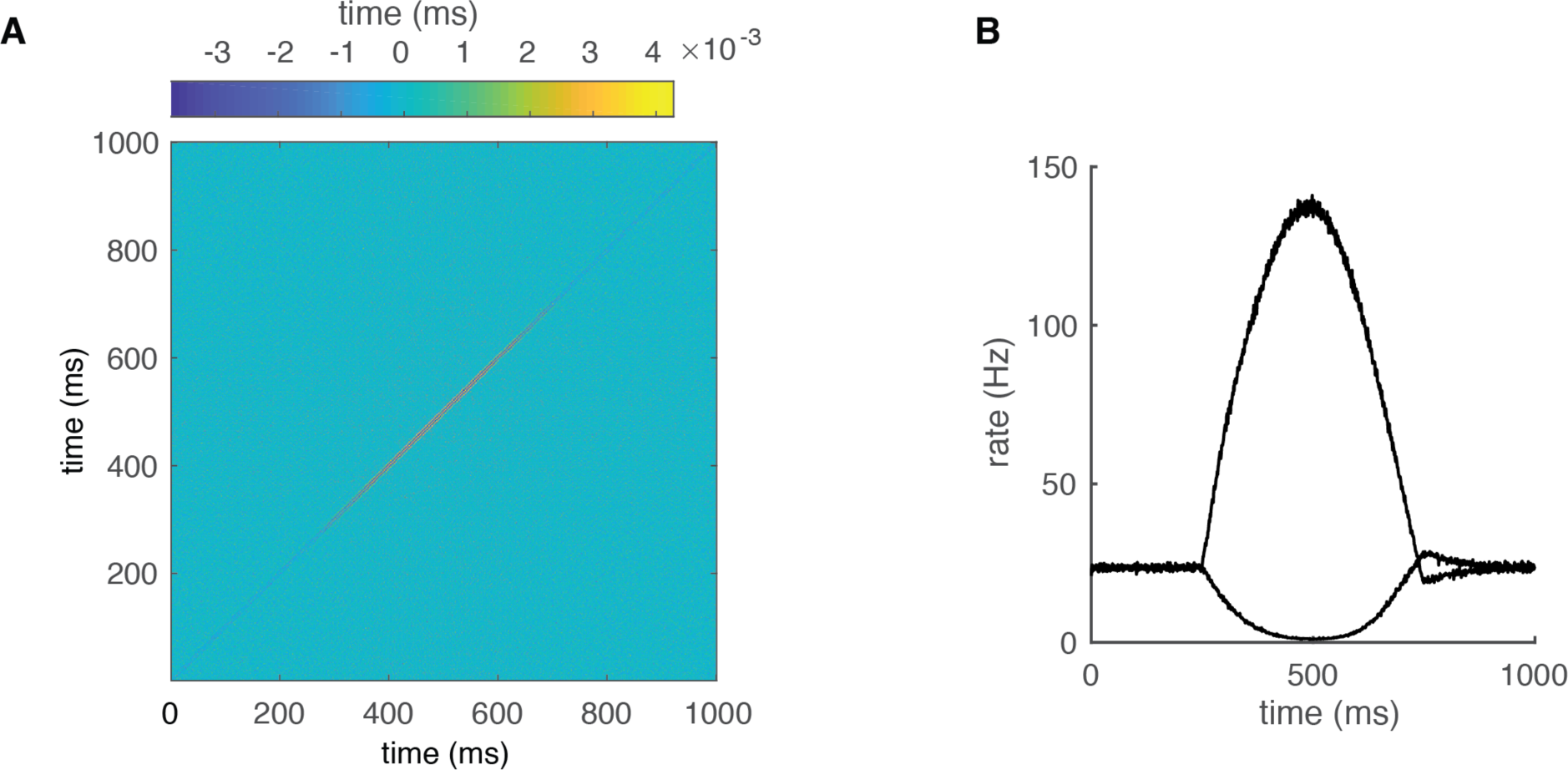
Dynamic Correlations of the PC Network Outputs (Correspond to Fig. 6). **A**. the JPSTH used to produce Fig. 6B. **B**, Population firing rates of increased-rate cells and decreased-rate cells for Fig. 6C-E.

**Figure S7.**
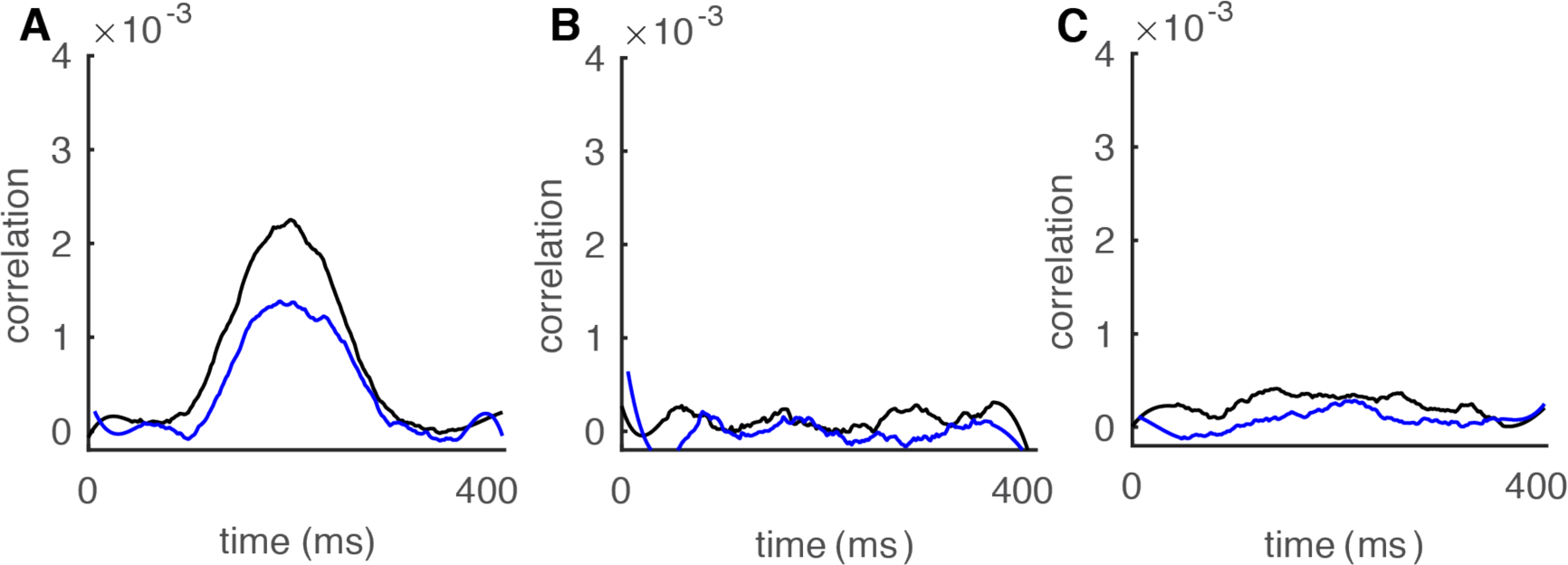
Transient Correlations for a 2.5 Hz sine wave. **A**. The 0 ms-time lag (black) and 6 ms-time lag (blue) correlations between increased-rate neurons in the network increase with population rates (n = 100); **B**. Correlations between decreased-rate neurons in the network (n = 30). **C**. Correlations between increased-rate neurons and decreased-rate neurons (n = 30 for each group).

## References

Abeles, M. (1982). Local cortical circuits : an electrophysiological study (Berlin ; New York: Springer-Verlag).

Aertsen, A.M., Gerstein, G.L., Habib, M.K., and Palm, G. (1989). Dynamics of neuronal firing correlation: modulation of “effective connectivity”. J Neurophysiol 61, 900–917.

Akemann, W., and Knopfel, T. (2006). Interaction of Kv3 potassium channels and resurgent sodium current influences the rate of spontaneous firing of Purkinje neurons. J Neurosci 26, 4602–4612.

Bartos, M., Vida, I., Frotscher, M., Meyer, A., Monyer, H., Geiger, J.R., and Jonas, P. (2002). Fast synaptic inhibition promotes synchronized gamma oscillations in hippocampal interneuron networks. Proc Natl Acad Sci U S A 99, 13222–13227.

Bishop, G.A., and O’Donoghue, D.L. (1986). Heterogeneity in the pattern of distribution of the axonal collaterals of Purkinje cells in zone b of the cat’s vermis: an intracellular HRP study. J Comp Neurol 253, 483–499.

Brunel, N., and Hakim, V. (1999). Fast global oscillations in networks of integrate-and-fire neurons with low firing rates. Neural Comput 11, 1621–1671.

Brunel, N., and Wang, X.J. (2003). What determines the frequency of fast network oscillations with irregular neural discharges? I. Synaptic dynamics and excitation-inhibition balance. J Neurophysiol 90, 415–430.

Buzsaki, G., and Draguhn, A. (2004). Neuronal oscillations in cortical networks. Science 304, 1926–1929.

Carnevale, N.T., and Hines, M.L. (2006). The NEURON book (Cambridge, UK ; New York: Cambridge University Press).

Chen, S., Augustine, G.J., and Chadderton, P. (2016). The cerebellum linearly encodes whisker position during voluntary movement. Elife 5, e10509.

Cheron, G., Gall, D., Servais, L., Dan, B., Maex, R., and Schiffmann, S.N. (2004). Inactivation of calcium-binding protein genes induces 160 Hz oscillations in the cerebellar cortex of alert mice. J Neurosci 24, 434–441.

Couto, J., Linaro, D., De Schutter, E., and Giugliano, M. (2015). On the firing rate dependency of the phase response curve of rat Purkinje neurons in vitro. PLoS Comput Biol 11, e1004112.

De Schutter, E., and Bower, J.M. (1994). An active membrane model of the cerebellar Purkinje cell. I. Simulation of current clamps in slice. J Neurophysiol 71, 375–400.

De Schutter, E., and Steuber, V. (2009). Patterns and pauses in Purkinje cell simple spike trains: experiments, modeling and theory. Neuroscience 162, 816–826.

de Solages, C., Szapiro, G., Brunel, N., Hakim, V., Isope, P., Buisseret, P., Rousseau, C., Barbour, B., and Lena, C. (2008). High-frequency organization and synchrony of activity in the purkinje cell layer of the cerebellum. Neuron 58, 775–788.

Destexhe, A., Rudolph, M., Fellous, J.M., and Sejnowski, T.J. (2001). Fluctuating synaptic conductances recreate in vivo-like activity in neocortical neurons. Neuroscience 107, 13–24.

Ermentrout, B., Pascal, M., and Gutkin, B. (2001). The effects of spike frequency adaptation and negative feedback on the synchronization of neural oscillators. Neural Comput 13, 1285–1310.

Ermentrout, G.B., Galan, R.F., and Urban, N.N. (2008). Reliability, synchrony and noise. Trends Neurosci 31, 428–434.

Fisyunov, A., Tsintsadze, V., Min, R., Burnashev, N., and Lozovaya, N. (2006). Cannabinoids modulate the P-type high-voltage-activated calcium currents in purkinje neurons. J Neurophysiol 96, 1267–1277.

Gauck, V., and Jaeger, D. (2000). The control of rate and timing of spikes in the deep cerebellar nuclei by inhibition. J Neurosci 20, 3006–3016.

Gutkin, B.S., Ermentrout, G.B., and Reyes, A.D. (2005). Phase-response curves give the responses of neurons to transient inputs. J Neurophysiol 94, 1623–1635.

Han, K.S., Guo, C., Chen, C.H., Witter, L., Osorno, T., and Regehr, W.G. (2018). Ephaptic Coupling Promotes Synchronous Firing of Cerebellar Purkinje Cells. Neuron 100, 564–578 e563.

Hebb, D.O. (1949). The organization of behavior; a neuropsychological theory (New York,: Wiley).

Heck, D.H., Thach, W.T., and Keating, J.G. (2007). On-beam synchrony in the cerebellum as the mechanism for the timing and coordination of movement. P Natl Acad Sci USA 104, 7658–7663.

Heiney, S.A., Kim, J., Augustine, G.J., and Medina, J.F. (2014). Precise control of movement kinematics by optogenetic inhibition of Purkinje cell activity. J Neurosci 34, 2321–2330.

Herzfeld, D.J., Kojima, Y., Soetedjo, R., and Shadmehr, R. (2015). Encoding of action by the Purkinje cells of the cerebellum. Nature 526, 439–442.

Hong, S., Negrello, M., Junker, M., Smilgin, A., Thier, P., and De Schutter, E. (2016). Multiplexed coding by cerebellar Purkinje neurons. Elife 5.

Khaliq, Z.M., Gouwens, N.W., and Raman, I.M. (2003). The contribution of resurgent sodium current to high-frequency firing in Purkinje neurons: an experimental and modeling study. J Neurosci 23, 4899–4912.

Kohn, A., and Smith, M.A. (2005). Stimulus dependence of neuronal correlation in primary visual cortex of the macaque. J Neurosci 25, 3661–3673.

Kuramoto, Y. (1984). Chemical oscillations, waves, and turbulence (Berlin ; New York: Springer-Verlag).

Lee, K.H., Mathews, P.J., Reeves, A.M., Choe, K.Y., Jami, S.A., Serrano, R.E., and Otis, T.S. (2015). Circuit mechanisms underlying motor memory formation in the cerebellum. Neuron 86, 529–540.

Maex, R., and De Schutter, E. (2003). Resonant synchronization in heterogeneous networks of inhibitory neurons. J Neurosci 23, 10503–10514.

Markowitz, D.A., Collman, F., Brody, C.D., Hopfield, J.J., and Tank, D.W. (2008). Rate-specific synchrony: Using noisy oscillations to detect equally active neurons. P Natl Acad Sci USA 105, 8422–8427.

Martina, M., Yao, G.L., and Bean, B.P. (2003). Properties and functional role of voltage-dependent potassium channels in dendrites of rat cerebellar Purkinje neurons. J Neurosci 23, 5698–5707.

Middleton, S.J., Racca, C., Cunningham, M.O., Traub, R.D., Monyer, H., Knopfel, T., Schofield, I.S., Jenkins, A., and Whittington, M.A. (2008). High-frequency network oscillations in cerebellar cortex. Neuron 58, 763–774.

Oemisch, M., Westendorff, S., Everling, S., and Womelsdorf, T. (2015). Interareal Spike-Train Correlations of Anterior Cingulate and Dorsal Prefrontal Cortex during Attention Shifts. J Neurosci 35, 13076–13089.

Orduz, D., and Llano, I. (2007). Recurrent axon collaterals underlie facilitating synapses between cerebellar Purkinje cells. Proc Natl Acad Sci U S A 104, 17831–17836.

Person, A.L., and Raman, I.M. (2012). Purkinje neuron synchrony elicits time-locked spiking in the cerebellar nuclei. Nature 481, 502–505.

Phoka, E., Cuntz, H., Roth, A., and Hausser, M. (2010). A new approach for determining phase response curves reveals that Purkinje cells can act as perfect integrators. PLoS Comput Biol 6, e1000768.

Rancz, E.A., and Hausser, M. (2010). Dendritic spikes mediate negative synaptic gain control in cerebellar Purkinje cells. Proc Natl Acad Sci U S A 107, 22284–22289.

Shin, S.L., and De Schutter, E. (2006). Dynamic synchronization of Purkinje cell simple spikes. J Neurophysiol 96, 3485–3491.

Singer, W. (1993). Synchronization of cortical activity and its putative role in information processing and learning. Annu Rev Physiol 55, 349–374.

Smeal, R.M., Ermentrout, G.B., and White, J.A. (2010). Phase-response curves and synchronized neural networks. Philos Trans R Soc Lond B Biol Sci 365, 2407–2422.

Traub, R.D., Jefferys, J.G.R., and Whittington, M.A. (1999). Fast oscillations in cortical circuits (Cambridge, Mass.: MIT Press).

Tsubo, Y., Takada, M., Reyes, A.D., and Fukai, T. (2007). Layer and frequency dependencies of phase response properties of pyramidal neurons in rat motor cortex. Eur J Neurosci 25, 3429–3441.

Uusisaari, M., and De Schutter, E. (2011). The mysterious microcircuitry of the cerebellar nuclei. J Physiol 589, 3441–3457.

van Welie, I., Roth, A., Ho, S.S., Komai, S., and Hausser, M. (2016). Conditional Spike Transmission Mediated by Electrical Coupling Ensures Millisecond Precision-Correlated Activity among Interneurons In Vivo. Neuron 90, 810–823.

Walter, J.T., Alvina, K., Womack, M.D., Chevez, C., and Khodakhah, K. (2006). Decreases in the precision of Purkinje cell pacemaking cause cerebellar dysfunction and ataxia. Nat Neurosci 9, 389–397.

Watt, A.J., Cuntz, H., Mori, M., Nusser, Z., Sjostrom, P.J., and Hausser, M. (2009). Traveling waves in developing cerebellar cortex mediated by asymmetrical Purkinje cell connectivity. Nat Neurosci 12, 463–473.

Watts, D.J., and Strogatz, S.H. (1998). Collective dynamics of ‘small-world’ networks. Nature 393, 440–442.

Witter, L., Rudolph, S., Pressler, R.T., Lahlaf, S.I., and Regehr, W.G. (2016). Purkinje Cell Collaterals Enable Output Signals from the Cerebellar Cortex to Feed Back to Purkinje Cells and Interneurons. Neuron 91, 312–319.

Zang, Y., Dieudonne, S., and De Schutter, E. (2018). Voltage- and Branch-Specific Climbing Fiber Responses in Purkinje Cells. Cell Rep 24, 1536–1549.

